# Rapid SARS-CoV-2 Detection and Classification Using Phase Imaging with Computational Specificity

**DOI:** 10.1101/2020.12.14.422601

**Authors:** Neha Goswami, Yuchen R. He, Yu-Heng Deng, Chamteut Oh, Nahil Sobh, Enrique Valera, Rashid Bashir, Nahed Ismail, Hyun J. Kong, Thanh H. Nguyen, Catherine Best-Popescu, Gabriel Popescu

## Abstract

Efforts to mitigate the COVID-19 crisis revealed that fast, accurate, and scalable testing is crucial for curbing the current impact and that of future pandemics. We propose an optical method for directly imaging unlabeled viral particles and using deep learning for detection and classification. An ultrasensitive interferometric method was used to image four virus types with nanoscale optical pathlength sensitivity. Pairing these data with fluorescence images for ground truth, we trained semantic segmentation models based on U-Net, a particular type of convolutional neural network. The trained network was applied to classify the viruses from the interferometric images only, containing simultaneously SARS-CoV-2, H1N1 (influenza-A), HAdV (adenovirus), and ZIKV (Zika). Remarkably, due to the nanoscale sensitivity in the input data, the neural network was able to identify SARS-CoV-2 vs. the other viruses with 96% accuracy. The inference time for each image is 60 ms, on a common graphic processing unit. This approach of directly imaging unlabeled viral particles may provide an extremely fast test, of less than a minute per patient. As the imaging instrument operates on regular glass slides, we envision this method as potentially testing on patient breath condensates.

The necessary high throughput can be achieved by translating concepts from digital pathology, where a microscope can scan hundreds of slides automatically.

**One Sentence Summary:** This work proposes a rapid (<1 min.), label-free testing method for SARS-CoV-2 detection, using quantitative phase imaging and deep learning.

## Introduction

COVID-19 is an infectious disease caused by the severe acute respiratory syndrome coronavirus 2 (SARS-CoV-2), which reached pandemic proportions in 2020. (*1*) The global impact of the disease on the healthcare systems and its socio-economic ramifications are severe and, likely, long-lasting.(*2*) The prompt response and public health measures have proven effective in limiting the spread of the virus, decreasing the number of active cases, and, ultimately the mortality rate. (*3*) Fast, accurate, and scalable testing has been recognized unanimously as crucial for mitigating the impact of COVID-19 and future pandemics. (*4*)

Diagnostic test accuracy is characterized by the sensitivity, defined as the probability of a positive result in a diseased patient, and specificity, given by the probability of a negative result in a healthy patient. Furthermore, the negative predictive value represents the chance of an individual with a negative test to be disease-free and, conversely, the positive predictive value is the chance that a person with a positive test is infected. In addition to these accuracy metrics, throughput and cost are important for deploying testing at scale. Recently, Weissleder et al. have reviewed the current status of the COVID-19 diagnostic tests (*4*). Briefly, nucleic acid tests (NATs) rely on the viral RNA being amplified via polymerize chain reaction (PCR) and are the most broadly used in the clinic today. NATs have been implemented on automated instruments and provide a result in several hours. Their accuracy may vary, with false negative rates reported in the order of 30% (*4, 5*). Serological tests assess the patient’s response to the viral infection through proteins such as immunoglobulin G. The efficacy of these tests relies on prior knowledge about the patient’s immune status as well as potential previous exposures to other virus types. The accuracy of serological tests is very high when performed ~20 days after the infection or first symptoms, but may lead to high false negative rates for early patients and false positives for patients previously exposed to other viruses (*4*). Common antigen tests can be performed using nasopharyngeal swabs and yield results in less than one hour. These tests operate on detecting proteins associated with the SARS-CoV-2 virus (nucleocapsid or spike proteins) using lateral flow or enzyme-linked immunosorbent assay (ELISA) tests.

Recently, accelerated efforts have been devoted to developing alternative testing procedures. These alternative detection schemes involve the use of plasmonic biosensors (*6-8*), fluorescence imaging of labelled virus particles and detection through machine learning (*9*), microfluidic immunoassays coupled with fluorescence detections (*10*) etc. While these approaches represent advances in SARS-CoV-2 detection methodologies, they still require either labelling or addition of foreign particles/solutions for the detection of SARS-CoV-2.

Here, we present a new approach for SARS-CoV-2 detection, which relies on direct, label-free imaging of viral particles. We employed spatial light interference microscopy (SLIM), a highly sensitive interferometric method, to image viruses deposited on a glass slide. Although, individual viruses are below the diffraction limit of the microscope, the optical path length information retrieved by SLIM unravels the nanoscale distribution of the refractive index associated with the individual and aggregated viral particles. We paired these data with deep learning algorithms, specifically optimized for viral particle detection and classification. Using fluorescence markers for specific virus tagging, we retrieved “ground truth” data by imaging the same field of view with both SLIM and epi-fluorescence. To emulate a more realistic application environment, we synthesized datasets where different virus types were “digitally mixed” onto the same SLIM image for deep learning development and evaluation. Thus, in addition to SARS-CoV-2, we imaged H1N1, HAdV and ZIKV. While a situation where a patient is exposed simultaneously to these four viruses is highly unlikely, we wanted to test it as a challenging task for our method and evaluate the specificity of our deep learning model. Following the training process, we tested the convolutional neural network (CNN) on unseen samples, classifying one virus type vs. the rest. Our results indicated a 96% area under the receiver operating characteristic curve for SARS-CoV-2, 99% for H1N1, 92% for HAdV and 91% for ZIKV.

This pre-clinical study demonstrates that sensitive imaging of unlabeled particles, paired with artificial intelligence (AI) can provide the foundation for a rapid, high-throughput, scalable test. The fact that the assay can be performed on the specimen placed on a glass slide allows for simple and fast sample collection, via, e.g., breath condensates. The image acquisition and inference take 100 ms total, which means that the entire test, including specimen collection, can be performed within a minute. Throughput can be scaled-up by borrowing engineering concepts from whole slide scanners in digital pathology, where hundreds of slides can be automatically fed into the imaging instrument. As the specimen requires minimum preparation and the instrument can be made portable, in principle, the technology can be deployed as a point-of-care solution.

The paper is structured as follows. First, we present the workflow for multimodal imaging and ground truth data acquisition. Next, we describe the SLIM imaging system and its sensitivity to the nanoscale ultrastructure of viral particles. We show 3D tomograms of the four virus types, to illustrate the subtle texture difference that the instrument captures, which the AI tools exploit for classification. We describe the convolutional neural network, which is a version of U-Net optimized for this problem. Finally, we present the accuracy of classifying the four virus types. We end with a discussion of the next steps necessary to implement this technology as a reliable clinical testing solution.

## Results

### Workflow

Figure 1 depicts the workflow of our approach (see Fig. S1 and Supplementary Section S1 for details on sample preparation). We tagged the deactivated virus samples with Rhodamine B isothiocynate as detailed in Materials and Methods. The staining was followed by dialysis to remove unbound fluorophores. The sample was deposited on a glass slide, fixed with EtOH, and air dried (Fig. 1A). The slide was imaged using multimodal SLIM and epi-fluorescence, overlaid for the same field of view (Fig. 1B). The resulting images were processed to extract pairs of images associated with individual particles (Fig. 1C). A U-Net convolutional neural network was trained using these data, with the fluorescence images acting as ground truth. The U-Net output provides a semantic segmentation map, i.e., an image that classifies and labels the various virus types (Fig. 1D).

**Figure 1.**
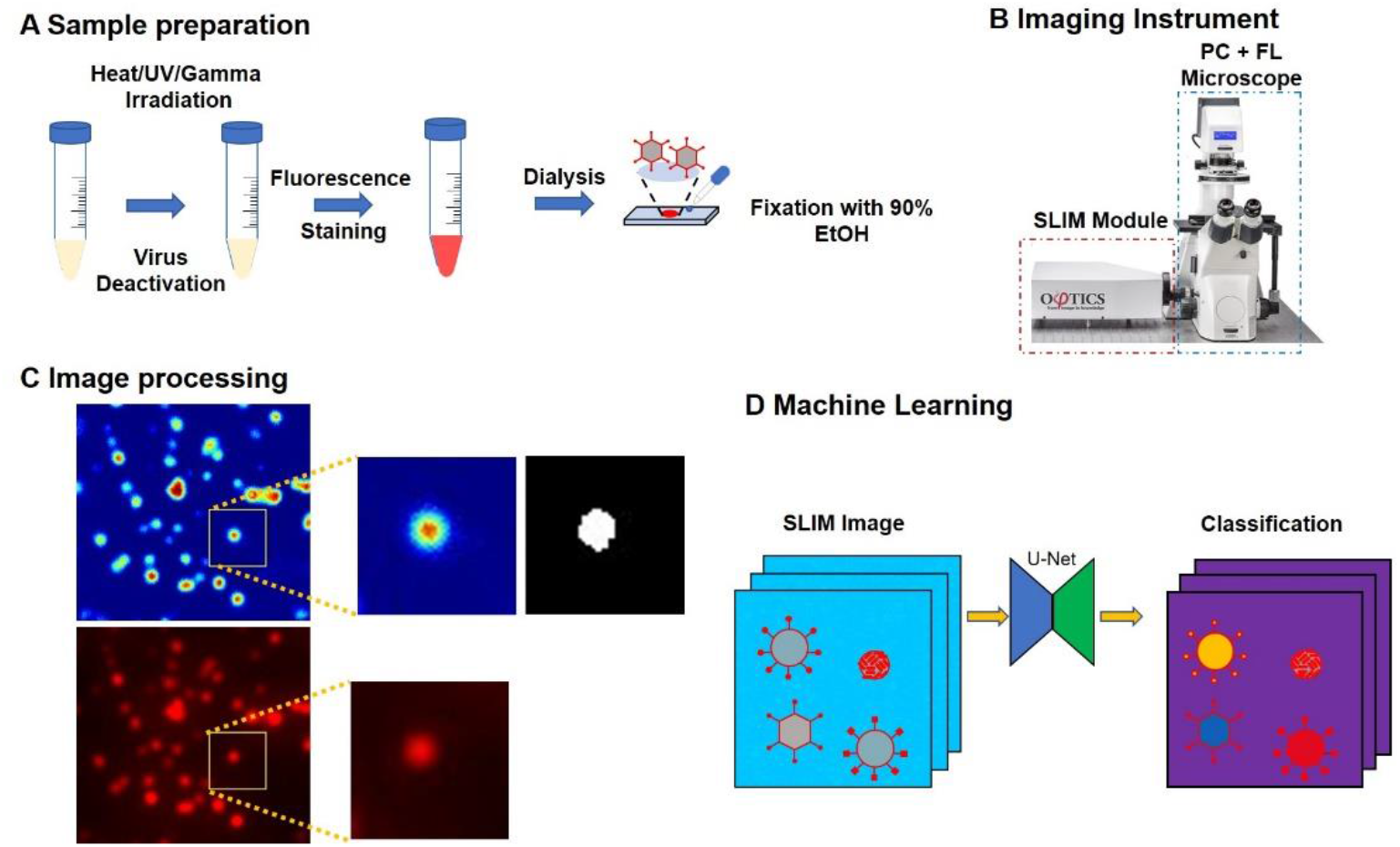
Virus particle classification using SLIM and machine learning. **A.** Sample preparation protocol, viruses were deactivated, stained with Rhodamine B isothiocyanate and dialyzed for 2 days to reduce fluorescence background and then placed on slide, fixed with 90% EtOH and air-dried **B.** We added a SLIM module to a traditional phase contrast microscope for quantitative phase information. **C.** SLIM and fluorescence were registered, single 48 x 48 regions were cropped from the image and segmented to provide label for multiclass classification. **D**. We synthesized a new dataset by randomly placing the cropped virus particles onto a background image acquired during the same experiment. A deep neural network was trained with this dataset to perform virus particle classification. Given a SLIM image, the model will output a class label for each pixel in the image.

### Imaging procedure

A key element in our approach is the spatial light interference microscope described in Fig. 2A. SLIM belongs to the family of quantitative phase imaging (QPI) instruments (*11*) which have found broad applications in biomedicine (*12-23*) due to their ability to image unlabeled, highly transparent structures. SLIM is implemented as an add-on module to an existing phase contrast microscope and, in essence, controls rigorously the phase shift between the incident and scattered field emerging from the specimen (*24, 25*). We used a Nikon Eclipse Ti inverted microscope outfitted with a SLIM module (CellVista SLIM Pro, Phi Optics, Inc.), which allows for fully automated data acquisition. The microscope objective pupil is relayed onto the surface of a phase-only spatial light modulator (SLM), such that the phase shift between the incident and scattered light is controlled precisely (Fig. 2A). We record four intensity frames associated with individual phase shifts, applied in increments of 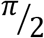, as shown in Fig. 2B. The four intensity images are combined as described in (*25, 26*) to decouple the amplitudes of the incident and the scattered fields from the phase information and obtain a quantitative phase map associated with specimen (Fig. 2B). Because the interfering fields in SLIM propagate along a common path, the phase measurement is highly stable, to within a fraction of a nanometer pathlength (*25*). Due to the white light illumination associated with the phase contrast microscope, the SLIM images are free of speckles, which converts into sub-nanometer spatial pathlength sensitivity (*25*). These attributes make SLIM ideal for the challenging task of imaging viral particles on a glass slide. Figure 2C illustrates the significant boost in contrast present in SLIM compared to traditional phase contrast microscopy.

**Figure 2.**
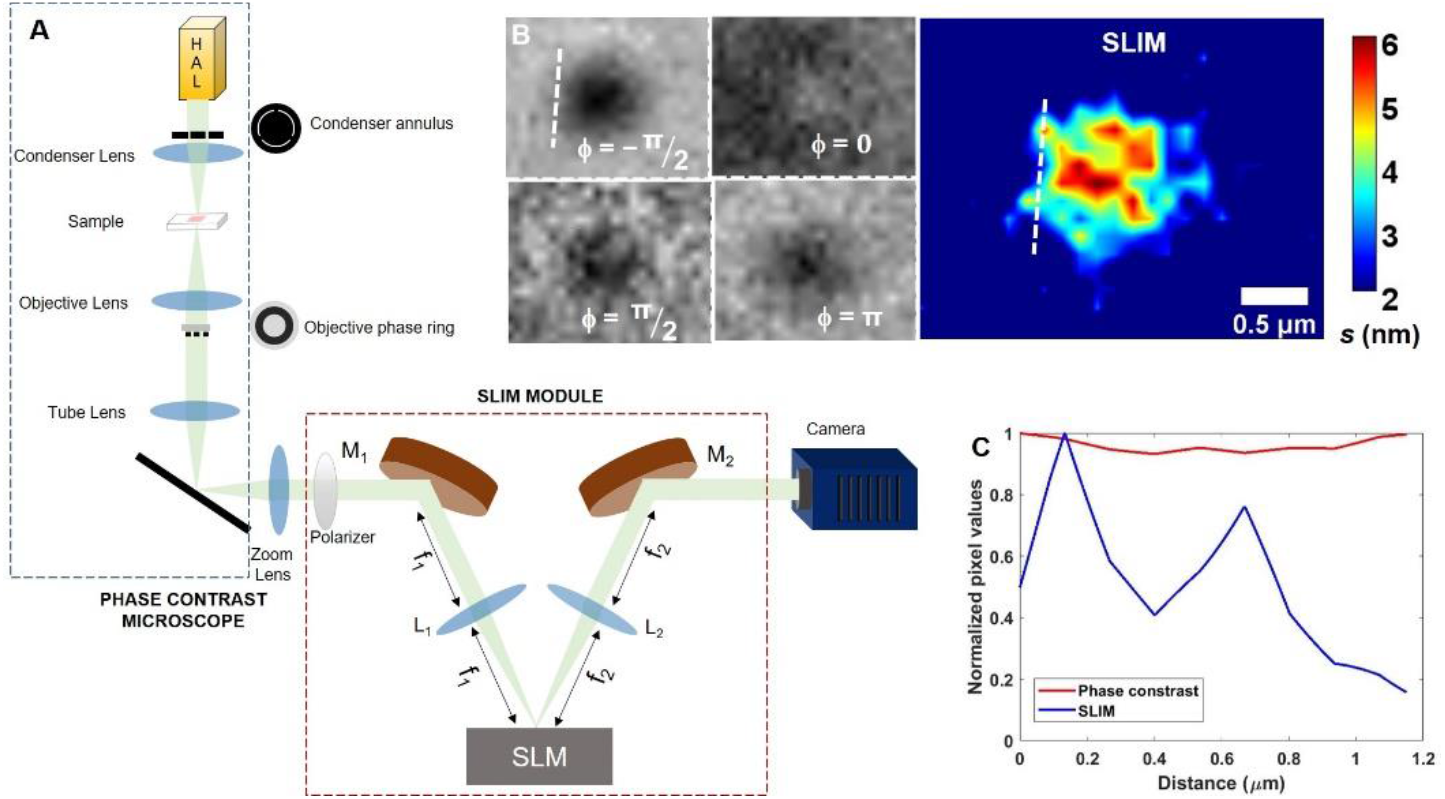
SLIM: **A.** Optical configuration of SLIM **B**. Image reconstruction, with color bar representing optical path length (s), in nm **C.** Profile through the dotted line in B, showing high sensitivity of SLIM over phase contrast.

### Virus detection and classification via SLIM

SARS-CoV-2, H1N1, HAdV and ZIKV were separately stained as illustrated in Fig. S1 (see Methods Section and Supplementary Section S1 for more details) with Rhodamine B isothiocyanate which has an emission at 595nm. We performed dual channel phase-fluorescence imaging on the samples. Figure 3 illustrates the imaging results for SARS-CoV-2, with SLIM (Fig. 3A) and fluorescence (Fig. 3B) images obtained on the same field of view. We registered the dual channel images using MATLAB for perfect overlay (see Supplementary Section S2 for details on image acquisition and processing). The regions denoted by the dash rectangular selections in Figs. 3(A, B) are zoomed-in and shown in Figs. 3(C, D). The discrete particles shown in the yellow rectangles reveal a 100% correspondence between phase and fluorescence, proving that SLIM is sensitive to the refractive index of the viral particles. For machine learning, we cropped out single particles within 48 × 48 pixel images. Figures 3 (E-J) show two examples of the cropped image set comprising of SLIM (Fig. 3(E, H)), fluorescence (Fig. 3(F, I)) and binary mask (Fig. 3(G, J)).

**Figure 3.**
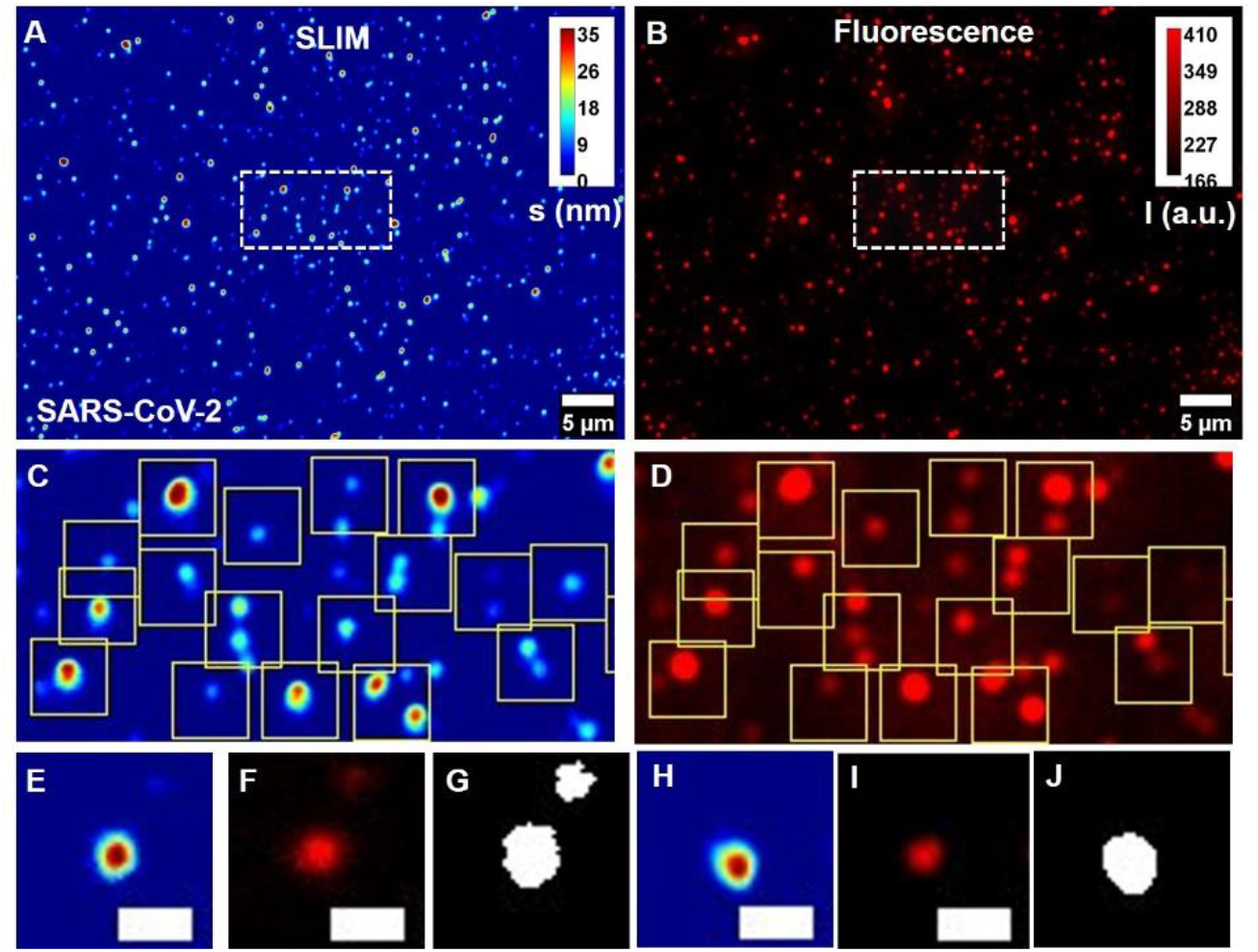
Correlated SLIM-Fluorescence imaging results for SARS-CoV-2: **A.** SLIM, colorbar represents optical path length fluctuations in nm, and **B.** fluorescence image, colorbar represents intensity in a.u., for the same field of view. **C, D.** Cropped SLIM and fluorescence images from the region inside white rectangle in A and B., yellow boxes highlight correspondence between SLIM and fluorescence. **E**. One 48 x 48 cropped image of SLIM, **F.** fluorescence and **G**. corresponding segmentation mask prepared for AI. Another cropped set for **H.** SLIM, **I.** fluorescence **J.** segmentation mask. Scale bar represents 5μm for A, B and 1μm for **E-J.**

Following same imaging procedure, we imaged H1N1 (Fig. S2), HAdV (Fig. S3) and ZIKV (Fig. S4) for SLIM and fluorescence. We cropped out 48 × 48 pixel images and performed segmentation to produce labels for four classes of virus. Although our images are still diffraction limited, SLIM’s nanoscale sensitivity to pathlength allows for efficient detection of viral particles.

The ultrastructure present in our SLIM data is demonstrated in deconvolved images (Supplementary Section S3). Using this operation, one can see that clumps of particles can be separated via deconvolution (Fig. S5).

### Deconvolution SLIM

Resolution of our imaging system is approximately 335nm (illumination at 550nm, objective 100x/1.45 with condenser NA 0.55). Following Rayleigh’s resolution criterion, two objects with separation less than the width of point spread function (PSF), cannot be fully resolved. The individual virus particles used in this study have an average diameter of less than 150nm, which makes them sub-diffraction objects for optical imaging. In order to push the resolution beyond the diffraction limit, we performed a deconvolution with the microscope’s PSF (Supplementary Section S3). To estimate the PSF, we identified the smallest spot in the images via a Matlab script. Using this PSF, the images were deblurred by employing the iterative Richardson-Lucy algorithm with total variation regularization (see Supplementary Section S3 for more details) (*27, 28*). Figure S5 illustrates the deconvolution results for the four virus classes. Thus, the deconvolution is able to produce deblurred images with clumps separated into smaller groups. However, it should be noted that the size of the deconvolved particles does not necessarily match the actual size of the virus particles as the decoupling of PSF and virus is still not perfect. However, we can successfully separate clumps into subsequent individual viruses, which the neural network is likely to pick-up for classification.

### Quantitative analysis

One advantage of SLIM over fluorescence is the inherent ability to measure not only shape descriptors like, diameter, orientation, circularity etc., but also quantify the phase information associated with the sample, which can then be used to extract biophysical information, such as, cell dry mass density. From the SLIM images, we extracted the total dry mass and surface dry mass density for each measured particle (see Supplementary Section S3 for details). We observed shifts in the dry mass density for different virus classes as shown in Fig. S6A. Figure S6(B-D) with p-values 1.35e-12, 8.84e-6 and 1.23e-5, respectively, demonstrate the statistical significance of the dry mass density differences between SARS-CoV-2 and H1N1, HAdV and ZIKV respectively, obtained by applying Kruskal-Wallis test (in MATLAB). These results indicate that dry mass density, which is incorporated in the SLIM data, is a marker that helps the machine learning algorithm to detect SARS-CoV-2.

### Tomographic Reconstructions

To get a better understanding of the viral particles, we performed a tomographic reconstruction of diffraction limited SLIM, using the Amira (Thermo Scientific) software (see Supplementary Section S4 for details). The results are shown in Fig. 4, where volumetric reconstructions of the particle cores (Fig. 4 (A-D)), and surface reconstructions (Fig. 4 (E-H)) for each particle are illustrated. These reconstructions provide an insight into structural dissimilarities that exist even in the diffraction limited SLIM images. Surface irregularities can be seen for SARS-CoV-2 in Fig. 4 (A, E). Figure 4 (B, F) show the H1N1 particle, which again has irregular surface but of different texture. Figure 4 (C, G) show a clump of at least two HAdV particles with hexagonal boundary visible in lower portion of Fig. 4G. ZIKV (Fig. 4 (D, H)) is significantly smoother compared to SARS-CoV-2. The structural signatures present in these reconstructions agree with the TEM images showing irregular surface morphology for SARS-CoV-2 (*29, 30*) and H1N1 (*37*), hexagonal cross-section for HAdV (*32*) and comparatively smoother surface of ZIKV (*33, 34*). These reconstructions suggest that signatures of structural information still exist in the diffraction limited SLIM images, due to the nanoscale pathlength sensitivity of SLIM. These subtle features help the machine learning algorithm to successfully classify these particles.

**Figure 4:**
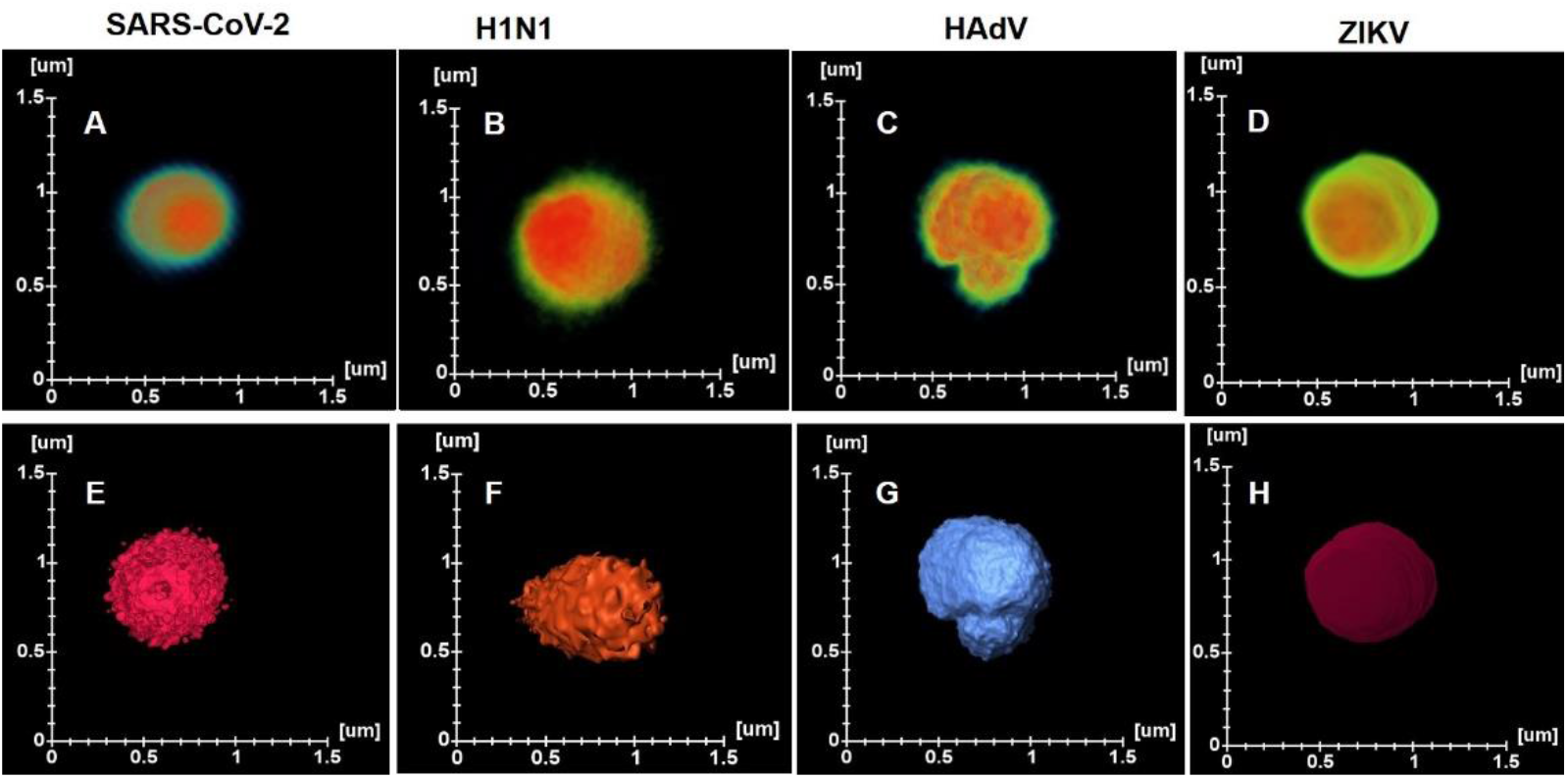
3D Tomograms: **Volume reconstruction of A.** SARS-CoV-2 **B.** H1N1 **C.** HAdV **D.** ZIKV. **Surface reconstructions of E.** SARS-CoV-2 **F.** H1N1 **G.** HAdV **H.** ZIKV. All reconstructions were performed using the Amira Software.

To assess the structural differences on a large scale, we performed volumetric reconstructions of groups of particles. Supplementary movies S1-S4 show the overall structural differences in diffraction-limited SLIM images. It can be seen that the maximum intensity projections of four virus classes exhibit differences in the structure, mainly, irregular surfaces for SARS-CoV-2 (Supplementary movie S1) and H1N1 (Supplementary movie S2), while hexagonal projections for HAdV (Supplementary movie S3) and comparatively smoother surface for ZIKV particles (Supplementary movie S4).

### Convolution Neural Network

We formulated the virus detection task as a semantic segmentation problem: given an input SLIM image containing several virus particles, our model predicts a probability distribution for each pixel, denoting the chance of this pixel belonging to one of the 5 classes: background, SARS-CoV-2, H1N1, HAdV, and ZIKV. An argmax operation turns the model output into a class label for each pixel. As all our raw SLIM images were of pure-culture virus particles, we synthesized a new dataset via “digital mixing” for machine learning development and evaluation (see Supplementary Section S5 for details).

The deep neural network we used was adapted from the U-Net (Fig. 5A and Fig. S7A) (*35*). Our model was trained using the digitally mixed SLIM images as input and the corresponding segmentation maps as ground truth (Fig. 5(B-C) and Fig. S7(B-C)). We divided machine learning task into two steps. Two types of datasets were prepared based on two data curation strategies. First dataset was semiautomatic, with manual cropping followed by automatic segmentation, fixed concentration of viruses per digitally mixed image and placement of virus particles on a grid with artificial phase background. Second dataset was fully automatic, with automatic segmentation followed by automatic cropping, varying (but balanced) concentration of viruses per digitally mixed image and random placement of virus particles on a blank image for digital mixing.

**Figure 5.**
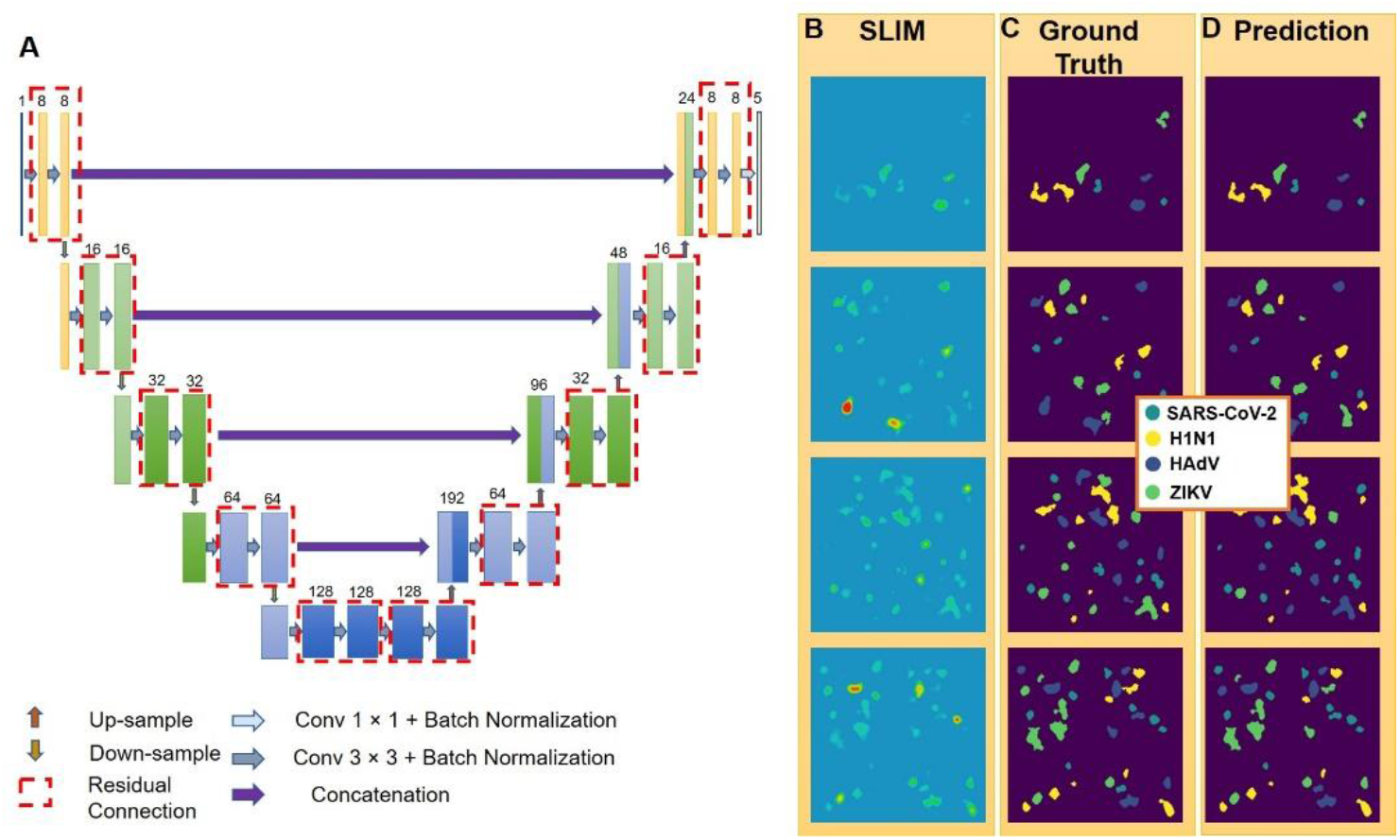
Training a deep neural network to perform classification of virus particles for the second dataset. **A.** We used a modified version of U-Net for this semantic segmentation task. Besides reducing the number of parameters in the network to around 0.8 million, we also added in residual connection and batch normalization for faster convergence. Model inference on images from the test set. **B.** Synthesized images of mixed virus particles. **C.** ground truth label. **D**. model inference.

Our first model (Fig. S7) was a proof-of-concept test-run. We manually cropped out 48 × 48 pixels regions of single virus particles from the images for all four viruses, collecting approximately 1200 cropped images. These cropped images were segmented, digitally mixed with an artificial background (see Methods Section and Supplementary Sections S2 and S5). Every digitally mixed image has five particles per class. We kept 500 particles out as the test dataset, and trained the neural network on the remaining particles (see Supplementary Section 5). During evaluation, we noticed that our model sometimes predicted more than one label per particle. To solve this issue, we used a post-processing strategy to enforce particle-level consistency in our model prediction (see Fig. S8 and Supplementary Section S5 for details on post-processing method). After the post-processing, we achieved the following area under the ROC curve (AUC) values for four viruses (Fig. S9A): 98% for SARS-CoV-2, 98% for H1N1, 96% for HAdV and 97% for ZIKV. The average precision and recall for this model are: 0.80 and 0.88 (SARS-CoV-2), 0.82 and 0.73 (H1N1), 0.88 and 0.78 (HAdV), 0.82 and 0.84 (ZIKV) (see Fig. S9B).

Our model’s excellent performance on this small, test-run dataset was the first step achieved in the direction of clinically usable, fast testing method. For the second phase of development, we moved on to a more realistic approach for data curation. To avoid bias in data selection and to focus on automation, we employed automatic processing to segment all the images and then crop out 48 × 48 particles from each image, based on the bounding box information of each particle, through a MATLAB script (see Supplementary Section S2). We emulated real life scenario where concentration and position of particles per sample can vary. So, each image in our digitally mixed dataset had between 2 to 8 particles of each virus type, resulting in between 8 to 32 virus particles in total. In this dataset, all 4 types of virus particles were randomly placed onto over 1600, 240 × 240 blank (background removed by segmentation) images (see Supplementary Sections S2 and S5 for more details of the procedure). We randomly selected around 1000 images for training and kept the remaining 564 SLIM images as the test dataset to evaluate our model. Similar as the first dataset, we enforced instance-level consistency on our model prediction via the same post-processing step (see Supplementary Section S5 and Fig. S8). Figure 5D shows the predictions after post processing. Quantitative results for this dataset are shown in Figure 6, where Fig. 6A shows the one-versus-all receiver operating characteristic (ROC) curve and Fig. 6B shows the complete confusion matrix to better illustrate our model’s sensitivity. AUC for all four virus classes is above 91%. We anticipate that, in clinical situations, the most challenging issue will be to detect the SARS-CoV-2 class alone, or, occasionally, distinguish it from the influenza virus (H1N1). The fact that the areas under the curve yield values of 96% and 99%, for SARS-CoV-2 and H1N1, respectively, is very encouraging. Average precision and recall values on the test dataset are: 0.80 and 0.85 (SARS-CoV-2), 0.98 and 0.99 (H1N1), 0.73 and 0.73 (HAdV), 0.74 and 0.63 (ZIKV) (Fig. 6B).

**Figure 6.**
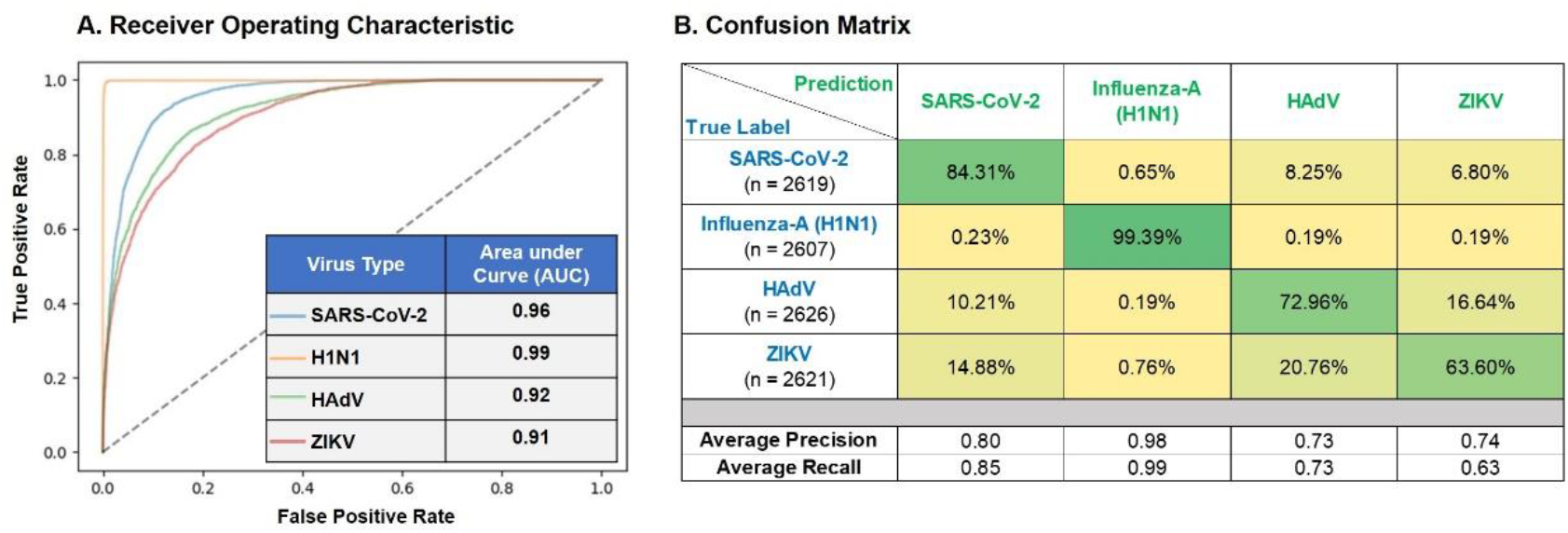
Model performance on the test dataset. **A.** The receiver operating characteristic (ROC) curve of the model on the test dataset. The model achieved over 0.9 area-under-curve (AUC) for all 4 virus types on the test dataset. The area-under-curve (AUC) for each class is computed by setting that class as label 1 and all other classes (the 3 remaining virus types and the background) as label 0. **B.** The confusion matrix of the model inference on the test dataset. Each row represents the ground truth label while each column represents the prediction. For visualization purposes, each entry in the confusion matrix was normalized with respect to the number of true labels (sum of each row). The precision, recall are averaged across all images in the test dataset. Both the ROC curve and the confusion matrix are evaluated on a per-particle level, where weighted average is computed to resolve conflict in model raw prediction.

We also plotted the precision and recall for SARS-CoV-2 on every image in the second test dataset into a histogram (Fig. S10). The majority of the detections have precision/recall values nearing unity. The learning curve plots for both our models (for first and second datasets) are shown in Fig. S11. The loss on the validation dataset and on the training dataset converged properly, indicating that our models did not overfit or underfit.

## Summary and Discussion

We presented a method for detection and classification of SARS-CoV-2 in the presence of other viruses, by using interferometric imaging and AI. Our results indicate that highly sensitive phase imaging is capable of providing subtle structural specificity of the viral particles, which in turn, allows for their accurate classification. There are two main components that help our model detect and classify viruses with high accuracy. First, the specific texture of the dry mass density can report on the differences in the refractive index caused by the specific protein compositions of the virus. Second, the nanostructure signature of individual viruses, e.g., irregularities on the surface of SARS-CoV-2 and H1N1, hexagonal shapes in HAdV, and the smoother surface of ZIKV, are subtle features in the SLIM images, exploited by the neural network.

The most likely combination of multiple viruses is SARS-CoV-2 and H1N1, a situation which can pose a challenge for accurate testing. However, our model proved to be successful in detecting and differentiating SARS-CoV-2 and H1N1 with a one versus all AUC of 96% and 99%, respectively. Pending successful clinical testing of this approach, we anticipate that the instrument can be implemented into a portable device controlled by a laptop. As the inference per field of view takes 60 ms, it is likely that the test per specimen, sampling several fields of view, will complete in a few seconds. Due to the lack of labels or other reagents, the test itself is bound to be inexpensive. Finally, to scale up throughput, we envision translating automatic slide scanning engineering concept from digital pathology devices.

## Material and Methods

### Sample preparation

The viruses used in this study are: Heat-inactivated SARS-CoV-2 (ATCC^®^ VR-1986HK™), Influenza A virus (H1N1) (ATCC^®^ VR-1894™), Human adenovirus 2 (HAdV) (ATCC^®^ VR-846™), and Zika (BEI: Zika Virus, PRVABC59, Infected Cell Lysate, Gamma-Irradiated (NR-50547)). HAdV and H1N1 were deactivated by UV. For fluorescence imaging, each virus solution was stained with Rhodamine B isothiocynate, separately for each experiment. Dialysis was carried out to remove unbound fluorophores from the stained solution. Stained virus sample was dropped on glass slide, fixed with 90% ethyl alcohol and air dried (more information in Supplementary Section S1).

### Image acquisition and processing

We performed dual channel correlative SLIM-fluorescence imaging on Nikon Eclipse Ti inverted microscope with add-on SLIM module (CellVista, Phi Optics, Inc.). Images were acquired with Nikon Plan-Apo 100x/1.45, phase contrast oil objective. Exposure was kept at 30ms and 200ms for SLIM and fluorescence, respectively. For 3D reconstructions, we acquired a z-scan passing through focus, with a step size of 5nm for the SLIM channel only. After the image acquisition, offline processing involved image registration of SLIM and fluorescence through MATLAB (see Supplementary Section S2). For the first dataset, we extracted 48 × 48 crops from SLIM and fluorescence images. We then segmented SLIM images to prepare the masks, which served as labels for the corresponding virus type during automated classification. For the second dataset, we first segmented the SLIM and fluorescence images and then performed automatic cropping based on bounding box information (more information in Supplementary Section S2).

We performed deconvolution using Richardson-Lucy iterative algorithm with Total Variation (TV) regularization (*27, 28*). We first converted phase map obtained from SLIM to complex field. This complex field was then used as an input to the algorithm. We derived an initial estimate for PSF from the images themselves, by choosing the smallest spot in the images. Utilizing the properties obtained from segmentation (area, integrated phase values, centroid, etc.,) we carried out quantitative analysis on single virus particles using MATLAB (see Supplementary Section S3).

We produced tomographic reconstructions using Amira software (Thermo Scientific). We cropped out single particles from whole image and upsampled them by a factor of 10 with bilinear interpolation to remove pixelations. We then used Volren and Isosurface rendering to reconstruct volume and surface tomograms (see Supplementary Section S4) for each virus type.

## Machine learning

For both the first (manual selection, with background) and second (automatic selection, without background) datasets, we prepared digitally mixed images to train and test our network. We placed single cropped viruses from each class, randomly in a 240 × 240 image, in fixed concentration for first dataset (5 particles per class) and varying concentrations (2 to 8 particles per class per image) for second dataset. During training, the model weights were updated using the Adam optimizer (*36*) against a categorical cross-entropy loss function. During evaluation, we found that in some cases, our model inferred more than 1 label for different parts of the same particle. To enforce instance-level consistency onto our model prediction, we performed a postprocessing step via connected component analysis to ensure that all pixels in each individual particle are predicted as one class. After this post-processing step (see Supplementary Section S5), our model’s performance was summarized into a confusion matrix on over 10,000 virus particles from the test dataset for the second dataset.

## Supporting information

Supplementary Movie S1. Volumetric reconstruction of a group of SARS-CoV-2 particles

Supplementary Movie S2. Volumetric reconstruction of a group of H1N1 particles

Supplementary Movie S3. Volumetric reconstruction of a group of HAdV particles

Supplementary Movie S4. Volumetric reconstruction of a group of ZIKV particles

## Acknowledgements

Heat-inactivated SARS-CoV-2 (ATCC^®^ VR-1986HK ™) was deposited by the Centers for Disease Control and Prevention and obtained through BEI Resources, NIAID, NIH: Genomic RNA from SARS-Related Coronavirus 2, Isolate USA-WA1/2020, NR-52285. Zika (BEI: Zika Virus, PRVABC59, Infected Cell Lysate, Gamma-Irradiated (NR-50547)) was obtained through BEI Resources, NIAID, NIH: Zika Virus, PRVABC59, Infected Cell Lysate, Gamma-Irradiated, NR-50547. Authors would also like to thank Dr. Mikhail E. Kandel and Dr. Catalin Chiritescu for providing the acquisition software CellVista (Phi Optics, Inc.) for SLIM control and dual channel imaging.

## Author contributions

G.P., C. B-P. and T.H.N. conceptualized and proposed the project. C. B-P., T.H.N., R.B., E.V. and C.O. carried out virus deactivation. H.J.K. and Y-H.D. selected chemical reagent appropriate for staining and carried out staining process on all four virus samples for fluorescence detections. N.G. prepared samples on slides, conducted imaging experiments, data analysis, tomographic reconstructions, deconvolution and data curation for machine learning. Y.R.H. and N.S. developed and trained machine learning model for virus detection and multiclass classification, performed post-processing and analysis of machine learning results. N.S. supervised the machine learning process. G.P., N.G. and Y.R.H. wrote the manuscript with contributions from all authors.

## Data and materials availability

All the data needed to reproduce the results can be obtained from the corresponding author upon reasonable request.

## List of supplementary materials

**Section S1.** Sample preparation

**Section S2.** Image acquisition and processing: registration, cropping and segmentation

**Section S3.** Deconvolution and quantitative analysis

**Section S4.** Tomographic reconstructions

**Section S5.** Machine learning: Development of deep learning models

**Figure S1: Fluorescence staining and sample preparation process:** The process of fluorescent tagging for virus particles.

**Figure S2. Correlated SLIM-Fluorescence imaging results for H1N1 Virus: A.** SLIM **B.** fluorescence for the same field of view, colorbar representing optical path length fluctuations in nm. **C, D.** Cropped single virus particles for SLIM and fluorescence, respectively. **E.** SLIM mask for AI training. Scale bar represents 5μm for A, B and 0.5 μm for C, D.

**Figure S3. Correlated SLIM-Fluorescence imaging results for HAdV: A.** Phase map obtained from SLIM, colorbar representing optical path length fluctuations in nm **B.** Fluorescence image for same field of view. **C** and **D** represent an example of one cropped virus particle (48 × 48 pixels) for SLIM and fluorescence respectively, with **E** representing the segmentation mask for labelling. Scalebar: 5μm for A, B and 0.5 μm for C, D.

**Figure S4. Correlated SLIM-Fluorescence imaging results for ZIKV: A**. SLIM **B**. fluorescence for the same field of view, colorbar representing optical path length fluctuations in nm. **C, D**. Cropped single virus particles for SLIM and fluorescence, respectively**. E.** SLIM mask for AI training. Scale bar represents 5μm for A, B and 0.5 μm C, D.

**Figure S5. Deconvolution results:** Inside each subfigure, raw SLIM images are in top row and deconvolved SLIM images are in bottom row for **A.** SARS-CoV-2 **B**. H1N1 virus C. HAdV, with hexagonal shape highlighted in 4th example and **D.** ZIKV, respectively. Scalebar is 0.5μm for all images.

**Figure S6. Quantitative analysis: A.** Dry mass density histogram for all four viruses, showing distinct peak for SARS-CoV-2. **B, C, D.** Kruskal-Wallis test results for dry mass density differentiation between SARS-CoV-2 and H1N1, HAdV and ZIKV, respectively. p-value in all cases is <0.0001, indicating high significance. Number of particles in each test is mentioned on the graphs.

**Figure S7. Training a deep neural network to perform classification on the first digital-mixing dataset. A.** We used a modified version of U-Net for this semantic segmentation task. Besides reducing the number of parameters in the network to around 3 million, we also added in residual connection and batch normalization for faster convergence. **Model inference on images from the validation set and the test set. B.** Synthesized images of mixed virus particles. **C.** ground truth label. **D.** model inference.

**Figure S8. Post-processing to enforce particle-level consistency. A.** To ensure all pixels in one virus particle has the same predicted label, we performed connected component analysis and averaged the probability distribution within each connected component. Left column: raw probability prediction; right column: probability distribution after post-processing. **B.** After postprocessing, the predicted segmentation map no longer had different labels within one particle-region. This enabled us to compute, on an instance-level, the performance of our model.

**Figure S9. Model performance on the first test dataset (consisting of 32 240 × 240 images). A.** The receiver operating characteristic (ROC) curve of the model on the test dataset. The model achieved over 0.96 area-under-curve (AUC) for all 4 virus types on the test dataset. The area-under-curve (AUC) for each class is computed by setting that class as label 1 and all other class labels (background and the 3 remaining virus types) as label 0. **B.** The confusion matrix of the model inference on the test dataset. Each row represents the ground truth label while each column represents the prediction. For visualization purposes, each entry in the confusion matrix was normalized with respect to the number of true labels (sum of each row). The precision, recall are averaged across all images in the test dataset. Both the ROC curve and the confusion matrix are evaluated on a per-particle level.

**Figure S10. Model Performance for SARS-CoV-2 with post-processing on the second dataset. A**. Histogram of particle-wise precision for SARS-CoV-2 evaluated on all 564 images in the test dataset. The average precision is 0.80. **B.** Histogram of particle-wise recall for SARS-CoV-2 evaluated on all 564 images in the test dataset. The average recall is 0.85.

**Figure S11. Learning Curve Plot. A.** The learning curve plot of our model developed for the first dataset. **B.** The learning curve plot of our model developed for the second dataset. Both plots showed a good convergence between the validation loss and training loss of our models, indicating that our models did not underfit or overfit. E represents categorical cross-entropy Loss

## References(37-62)

**Movie S1.** Volumetric reconstruction of a group of SARS-CoV-2 particles with maximum projection emphasizing the irregular boundaries of the virus particles.

**Movie S2.** Volumetric reconstruction of a group of H1N1 particles with maximum projection emphasizing the irregular boundaries of the virus particles and pleomorphic shapes of H1N1 particles.

**Movie S3.** Volumetric reconstruction of a group of HAdV particles with maximum projection emphasizing the hexagonal shape of the virus particles.

**Movie S4.** Volumetric reconstruction of a group of ZIKV particles with maximum projection emphasizing the relatively smoother surface of the virus particles.

