## Supplementary Text for "Rapid SARS-CoV-2 Detection and Classification Using Phase Imaging with Computational Specificity"

**S1. Sample preparation:**

**SARS-CoV-2**: Heat-inactivated SARS-CoV-2 was obtained from ATCC (ATCC® VR-1986HK™). The vial was centrifuged prior to opening. Since the virus was already heat deactivated, no further deactivation was performed. SARS-CoV-2 has a spherical structure with diameter ranging from 60-140 nm (*37, 38*). The surface has spike like protein protrusions (*38*).

**Human adenovirus type 2 (HAdV)** was purchased from ATCC (ATCC® VR-846™). A549 cell line was used as host for HAdV, which was also obtained from ATCC (CCL-185). In brief, HAdV was propagated in A549 cells supplemented by Ham F-12 media with 2% fetal bovine serum (Thermo Fisher Scientific, MA, USA), and 1X antibiotic-antimycotic (Thermo Fisher Scientific, MA, USA). After 5 days of incubation at 37°C with 5% $CO_2$ when 80% cytopathic effect was reached, HAdV was harvested from the cells by three freeze-thaw cycles. To purify HAdV, the virus solution was centrifuged at 2000 rpm (556 g) for 10 min (Sorvall Legend RT Plus, Thermo Fisher Scientific, MA, USA), and the virion-containing supernatant was filtered by a 0.45 μm membrane filter (Millipore Sigma, MA, USA). The virion-containing filtrate was then purified using ultracentrifuge with 36000 rpm (150700 g) at 4°C for 3 hours (Optima XPN-90 Ultracentrifuge, Beckman Coulter, CA, USA). The virus pellet on the ultracentrifuge tube was resuspended in 1X PBS and stored at -80°C before use. The infectivity of the purified HAdV was confirmed to be about $10^7$ PFU/mL by plaque assay.

The infectious HAdV was inactivated by ultraviolet (UV) irradiation at 254 nm. The UV irradiation at 254 nm was known to inactivate HAdV by primarily damaging genomic DNA (*39*). The UV irradiation at 254 nm was generated in this study using a medium-pressure UV generator (Calgon Carbon Co., Pittsburgh, PA) with a bandpass filter at 254 nm. We exposed 100 uL of virus solution to the UV irradiation (254 nm) for about 15 min which was equivalent to about 300 mJ/cm$^2$. According to Ref. (*39*), the fluence of 300 mJ/cm$^2$ was expected to inactivate HAdV by 10-log reduction. HAdV has an icosahedral shape, with diameter of about 100nm (*40*).

**Zika Virus (ZIKV):** The Zika virus, PRVABC59 (BEI: Zika Virus, PRVABC59, Infected Cell Lysate, Gamma-Irradiated (NR-50547)) was gamma-irradiated (5 x 106 RADs) on dry ice. The sample was diluted in Nuclease-free water. The ZIKV particles are spherical (approximately 50 nm in diameter) with a 30 nm electron-dense core (*41, 42*). The mature ZIKV contains 180 copies of the envelope protein (E) and membrane (M) proteins in icosahedral-like organization and arranged in the raft configuration (*41, 42*). The E protein predominates on the surface with the M protein residing underneath the E protein (*42*). The raft configuration consists of three E proteins dimers which lie parallel to one another, with the virion having a total of 30 rafts (*42*). ZIKV has an imperfect icosahedral structure with a diameter of approximately 50-60nm (*43, 44*).

**Influenza-A (H1N1)** was obtained from ATCC (ATCC® VR-1894™). UV irradiation with 254 nm wavelength was also applied to inactivate the viruses. Although UV irradiation could cause structural change in capsid proteins, the primary inactivation mechanism of the UV irradiation is known to mutagenize the genomes (*45*). The virus stock was used without neither further inoculation in cells nor the purification. The influenza virus was exposed to the same UV irradiation system described for HAdV inactivation, but the exposure time was 10 min which is equivalent to 200 mJ/cm$^2$. This fluence was expected to inactivate the viruses about 30 log-reduction assuming the inactivation rate follows the first-order reaction (*46, 47*). H1N1 is a spherically shaped particle with a diameter of approximately 100nm, it also features spike like protrusions on the surface (*48*). However, it may also exhibit pleomorphism resulting in elongated structure (*48*).

**Fluorescence Tagging**: All four virus particles were tagged separately with fluorescent probes in order to validate their presence for observation. For each experiment, virus particles were suspended in 5 mL of carbonate/bicarbonate buffer (0.1 M, pH=9.2). Then, 1 mL of rhodamine B isothiocyanate (RBITC) (2 mg/mL in DMSO) solution was added into the virus solution under stirring condition. The RBITC binds onto virus through conjugation between isothiocyante group and amine group on capsid of virus. The reaction was conducted for 2 hours and protected from light. In the end, the virus particles tagged with rhodamine B were purified using dialysis against deionized water. After 2 days, the purified virus particles tagged with rhodamine B solution was poured on glass slide for fixation with 90% Ethyl alcohol. Staining procedure is depicted in Fig S1.

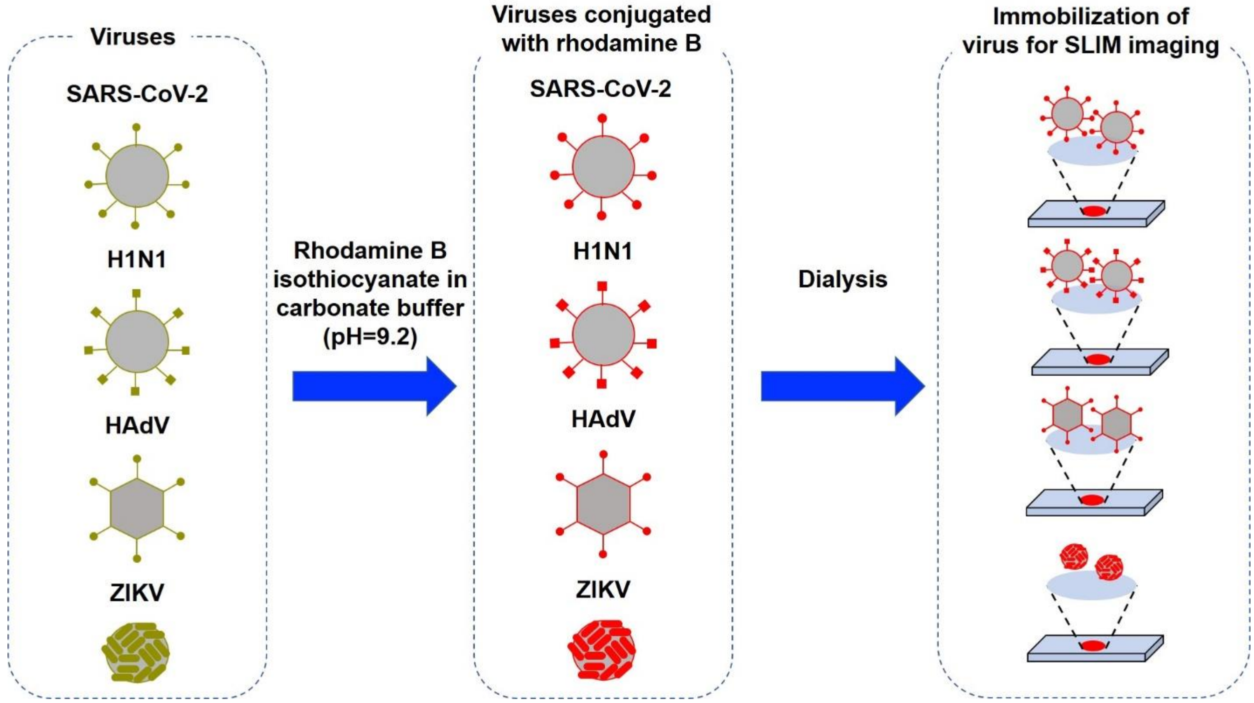

**Figure S1: Fluorescence staining and sample preparation process:** The process of fluorescent tagging for virus particles.

**Slide preparation:** 10µL of stained virus solution was dropped on a plain glass slide and allowed to air dry. Upon complete drying, the sample spot was treated with 90% Ethyl alcohol for fixation followed by air drying again. Completely dried sample was then covered with coverslip (#1) and the edges were sealed with nail polish.

### S2. Image acquisition and processing: registration, cropping and segmentation

**Image acquisition:** We performed dual channel phase-fluorescence imaging. Nikon Ti-E microscope was used for these experiments with SLIM module (CellVista SLIM Pro, Phi Optics, Inc.) connected to the left port of the microscope. SLIM acquisition was performed using Andor-Zyla camera. For the fluorescence, we used Zeiss AxioCam MRm camera mounted on the right port of the microscope. Imaging was done using Nikon Oil immersion 100x/1.45 phase contrast objective, with 18 pixels/µm resolution (including 1.2x zoom provided by SLIM module). For fluorescence, TRITC filter was used with exposure time 200ms. For each field of view, we used CellVista software (Phi Optics, Inc.) to capture a pair of SLIM and fluorescence images. SLIM can produce images at 15fps. Fluorescence was required for generating the ground-truth for machine learning on the phase data. For each field of view, synchronized and sequential acquisition was performed for the two channels. SLIM image was acquired on Andor-Zyla camera with a resolution of 1392 × 1040 pixels. Corresponding fluorescence images were captured by the Zeiss AxioCam MRm camera with an image size 1388 × 1040 pixels. Imaging results for SARS-CoV-2 are shown in the main text Fig. 3.

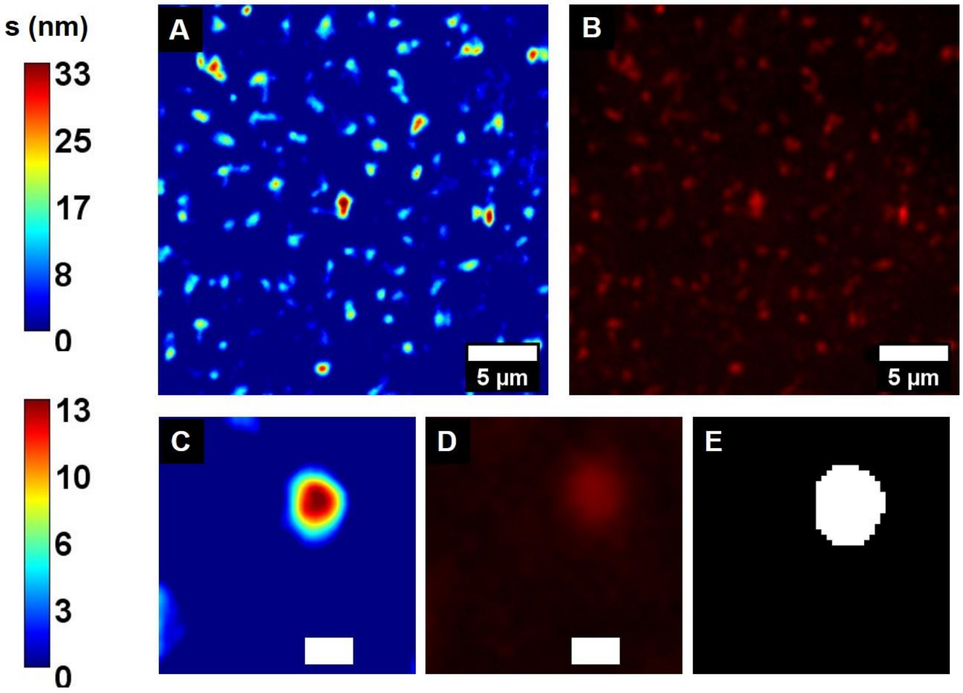

**Figure S2. Correlated SLIM-Fluorescence imaging results for H1N1 Virus: A.** SLIM **B.** fluorescence for the same field of view, colorbar representing optical path length fluctuations in nm. **C, D.** Cropped single virus particles for SLIM and fluorescence, respectively. **E.** SLIM mask for AI training. Scale bar represents 5µm for A, B and 0.5 µm for C, D.

Following the same procedure as outlined in main text, we imaged H1N1, HAdV and ZIKV. Figures S2-S4 represent the imaging results for H1N1, HAdV, and ZIKV, respectively. SLIM and fluorescence images for the same FOVs are shown in subfigures A and B, with 48 x 48 pixel

SLIM, fluorescence, and SLIM mask, respectively, shown in C, D and E.

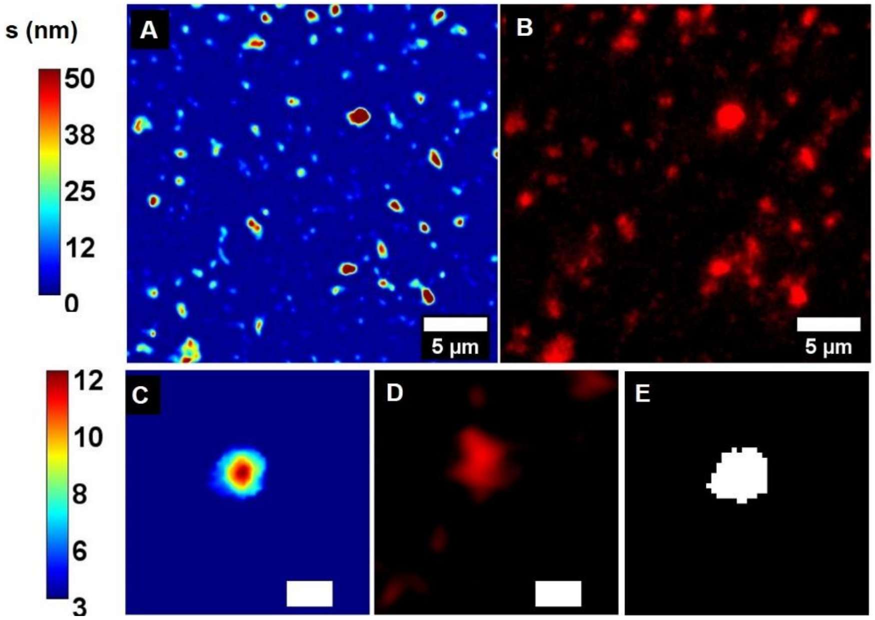

**Figure S3. Correlated SLIM-Fluorescence imaging results for HAdV: A.** Phase map obtained from SLIM, colorbar representing optical path length fluctuations in nm **B.** Fluorescence image for same field of view. **C** and **D** represent an example of one cropped virus particle (48 × 48 pixels) for SLIM and fluorescence respectively, with **E** representing the segmentation mask for labelling. Scalebar: 5µm for A, B and 0.5 µm for C, D.

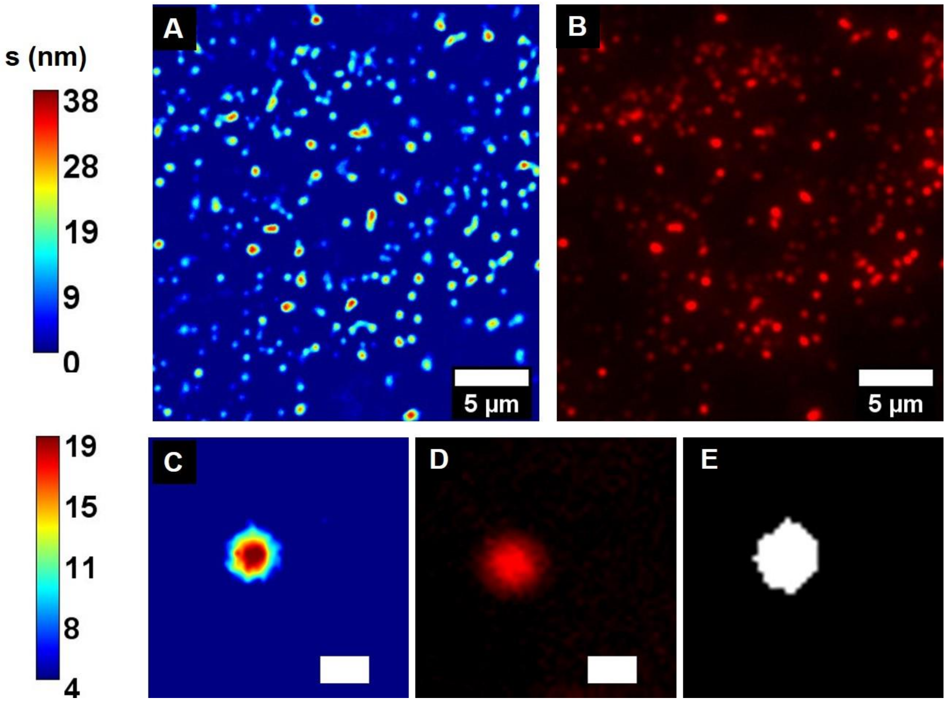

**Figure S4**. **Correlated SLIM-Fluorescence imaging results for ZIKV: A**. SLIM **B**. fluorescence for the same field of view, colorbar representing optical path length fluctuations in nm. **C, D**. Cropped single virus particles for SLIM and fluorescence, respectively**. E.** SLIM mask for AI training. Scale bar represents 5μm for A, B and 0.5 μm C, D.

Image overlay involved two sequential image registrations. MATLAB scripts were used for both the steps of registration. First registration was based on manual control point selection and similarity transformation. For the second step, the resultant transformed image was then used as a moving image in MATLAB app called Registration Estimator, using a 'multimodal' registration with translation.

For the first, test-run dataset, we manually selected 48 × 48 cropped images from both SLIM and fluorescence images using a macro written in FIJI (ImageJ) that synchronizes the coordinates of ROI selection for both images of same FOV. These cropped SLIM images were then segmented to be used as labels for machine learning algorithm. Care was taken to select single particle and avoid clumps.

For the second, main dataset, we prepared binary masks from the SLIM images. For training the neural network, 48 × 48 pixels cropped images were prepared based on bounding box information from 'regionprops' function (MATLAB). These masks would serve as a label for the four classes. We carried out segmentation using scripts written in MATLAB. Input images were median filtered with a neighborhood window of 5 pixels to reduce noise. Adaptive thresholding was used with a sensitivity of 0.65 and a neighborhood size of 17 pixels in each dimension. Morphological operations like open, fill and dilate were used to refine the mask and remove stray elements. Considering 335nm as our minimum detection spot width based on our system PSF, anything with area less than 29-pixel square was not selected in the final mask. Finally, to remove separate objects with borders touching, we used distance transformation and watershed algorithm. Same parameters were used to segment all four-virus dataset to avoid bias. For each detection, a label was created and 'regionprops' function (MATLAB) was used to measure all properties. These properties were used to extract sum of phase values over the area, equivalent diameter, centroid, circularity, bounding box coordinates and pixel index of the area to be used for quantitative analysis to calculate dry mass, dry mass density and selection of PSF estimate (through equivalent diameter and circularity). We imaged 13,143 SARS-CoV-2, 18,763 H1N1, 9,346 HAdV, and 12,299 ZIKV particles. The lower number of detections in HAdV can be attributed to the clumping nature that we observed in the sample.

We also prepared fluorescence masks to validate our SLIM selections. The segmentation procedure was the same as for SLIM. However, for the adaptive thresholding, we chose sensitivity to be 0.55, with a neighborhood size of 27 pixels along each dimension.

**S3. Deconvolution and quantitative analysis:** For the deconvolution, the smallest spot in the images was identified through a script in MATLAB and assumed to be the initial estimate of the PSF. Using this PSF estimate for deconvolution, the blurred image was restored. Deconvolution was performed in MATLAB. We used iterative, Richardson-Lucy (RL) algorithm in conjunction with total variation (TV) regularizer (*27, 28*).

The algorithm works on the complex field image, which is defined as

$$I = e^{j\phi(x,y)}, \qquad [1]$$

where ϕ(x,y) is the SLIM image. The algorithm iteratively solves for (*27, 28*)

$$I^{m+1}(x,y) = A(x,y)\left[ I_{psf}(-x,-y) \otimes \frac{I^{o}(x,y)}{I_{psf}(x,y) \otimes I^{m}(x,y)} \right] \qquad [2]$$

Where $I^0$ is the observed image, $I_{psf}$ is the estimate of PSF, and A(x, y) is the combination of the present image and Total Variation regularizer, given by (*27, 28*)

$$A(x,y) = \frac{I^{m}(x,y)}{1 - \beta\nabla.\left( \frac{\nabla I^{m}(x,y)}{\left\| I^{m}(x,y) \right\|} \right)} \qquad [3]$$

In Eq. 3, β is regularization parameter. The iteration is terminated when there is no significant change between iterations. Finally, the phase is extracted from last iteration results to yield deconvolved SLIM.

$$\hat{\phi}(x,y) = \arctan\left[ \frac{imag\left( I^{m+1}(x,y) \right)}{real\left( I^{m+1}(x,y) \right)} \right] \qquad [4]$$

Smallest detection in each dataset for all four-viruses was used as the initial estimate of PSF. To get this estimate, we selected particles with equivalent diameter less than 7 pixels (which translates to 388nm, to cover our estimated width of PSF, 335nm) and circularity greater than 0.8. Out of this selected set, we selected random samples of images based on visual feedback and applied 'Smooth' operation in FIJI (ImageJ) on these selected images to reduce spurious pixel noise. We generated surface plot for each of the selections and the images with multiple peaks were discarded. Final estimates were then tested on the deconvolution algorithm one by one to select the best estimate that would not introduce significant ringing, edge effects or noise. We note that although the deconvolution is not able to fully resolve the individual viruses, it still deblurs the images to show morphological evidence (hexagonal shape, in case of HAdV, Fig S5C) of the virus structure in the detected particles. Figure S5 shows the deconvolution results for all four viruses.

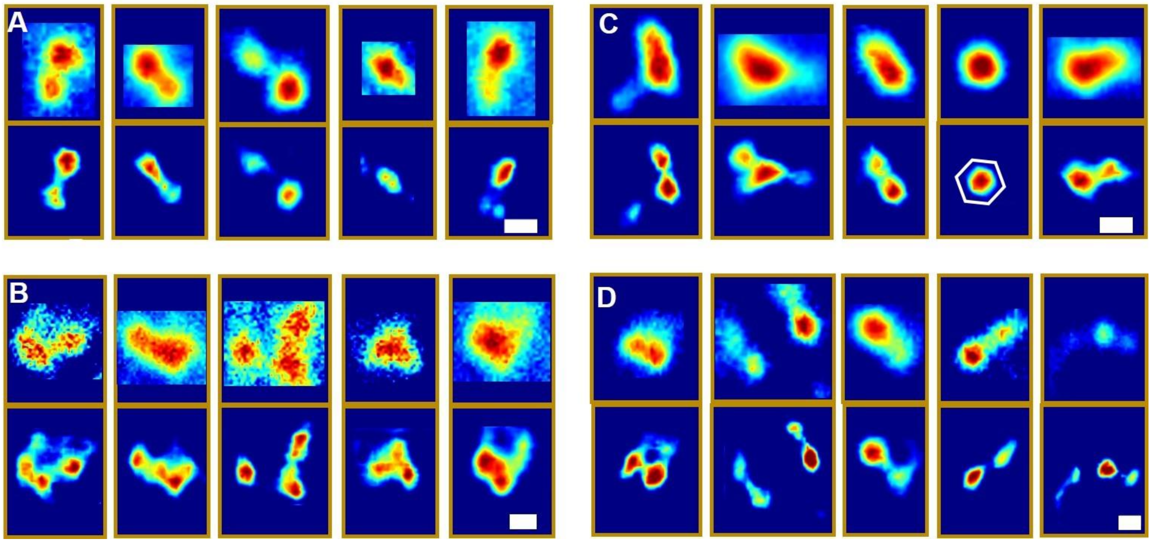

**Figure S5 Deconvolution results:** Inside each subfigure, raw SLIM images are in top row and deconvolved SLIM images are in bottom row for **A.** SARS-CoV-2 **B.** H1N1 virus **C.** HAdV, with hexagonal shape highlighted in 4$^{th}$ example and **D.** ZIKV respectively. Scalebar is 0.5μm for all images.

**Quantitative analysis: Dry mass, dry mass density**

Dry mass is a measure of non-aqueous mass of the biological sample (*11*). It is defined as *(11)*

$$M=\frac{\lambda}{2\pi\eta}\iint_A \phi(x,y)\mathrm{dxdy} \quad [5]$$

Dry mass density is the ratio of dry mass to area and is defined as

$$\rho(\mathrm{x,y})=\frac{\lambda}{2\pi\eta}\phi(\mathrm{x,y}) \quad [6]$$

In Eq. 6, λ is the central wavelength of illumination, 550nm, η is the refractive index increment with a value of 0.2mL/g (*11*) and ϕ is the phase shift introduced by the sample.

We calculated and compared dry mass density of carefully selected single virus images. Fig. S6A shows the histogram of dry mass density for all four viruses, which clearly shows the peak separations between the four classes. To investigate the statistical significance of these peak differences, we applied Kruskal-Wallis test (due to non-Normality of the data) in MATLAB on the dry mass densities of all four virus classes. Results are shown in Fig. S6 (B-D) for dry mass density difference between single virus particle images selected by MATLAB script for SARS-CoV-2 and H1N1, HAdV and ZIKV respectively. The p-values are 1.35e-12, 8.84e-6 and 1.23e-5 for Fig. S6 (B-D) respectively showing the high significance of dry mass density difference of SARS-CoV-2 and other viruses in this study.

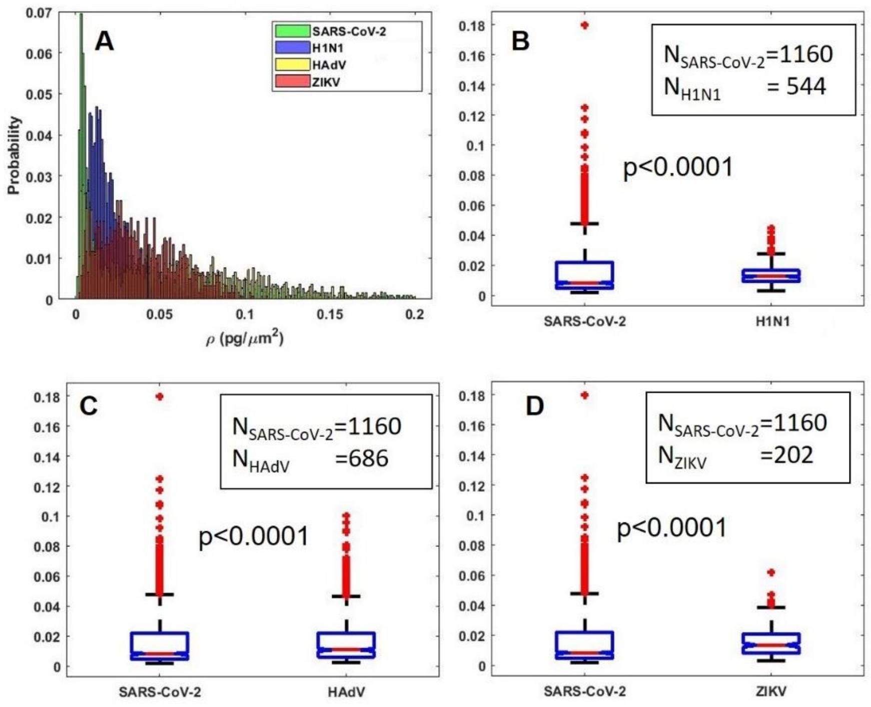

**Figure S6**. **Quantitative analysis: A.** Dry mass density histogram for all four viruses, showing distinct peak for SARS-CoV-2. **B, C, D.** Kruskal-Wallis test results for dry mass density differentiation between SARS-CoV-2 and H1N1, HAdV and ZIKV respectively. p-value in all cases is <0.0001, indicating high significance. Number of particles in each test is mentioned on the graphs.

### S4. Tomographic reconstructions

Motivated by the structural signatures provided by SLIM, we investigated the possibility of retrieving more information out of the 3D reconstructions from z-scan of SLIM images. For this purpose, we did a z-scan SLIM acquisition, ~2 μm above and below the focus of particles, with a step size of 1-5nm, using a 100x/1.45 NA objective. Out of the z-stack, substack covering slices extending from just above to just below the surface of the virus particles was selected. For the reconstruction, we used the Amira (Thermo Scientific) software and segmented images either through MATLAB or through Amira's functions, 'labelfield' and 'segmentation', depending on the accuracy of the resulting label. Images were resized by a factor of 10 and bilinear interpolation was used for better visualization. Volumetric rendering was performed in Amira, after a histogram equalization of intensity to enhance contrast. Isosurface rendering was performed to create surface rendering. Both the volumetric and surface reconstructions through Amira are shown in main text Fig. 4, and supplementary movies S1-S4.

### S5. Machine learning: Development of deep learning models

**Digital Mixing:** For the training and subsequent test and validation of our network, we employed a scheme called digital mixing. 48 × 48 cropped images of all four detected virus particles (SARS-CoV-2, H1N1, HAdV, and ZIKV) were mixed in fixed (n=5 per class per image, first dataset) and varying (n=2 to 8 per class per image, second dataset) proportions to emulate the situation where the four viruses were physically mixed on a slide. This operation was achieved using scripts written in MATLAB. For the second dataset, seven sets of images, each of 240 × 240 pixels, were generated. Each set contained 2 to 8 particles from each class, for a total of 1611 digitally mixed images. We used colocalized image pairs to train a deep convolutional neural network to map any phase image to a segmentation mask that provides a pixel-wise label for the background, SARS-CoV-2, H1N1, HAdV and ZIKV.

#### Model Architecture and Training Strategies

For the deep convolutional neural network, we picked a variant of U-Net (*35*) that has shown great performance on segmentation tasks with quantitative phase imaging (QPI) data in our previous works (*49*-*51*). The network contains an encoder path, a bottleneck connection and a decoder path (Fig. 5A). The encoder path is responsible for extracting features from the input image via 4 stages of building blocks containing convolution layers interlaced by downsampling operations. Unlike the original U-Net, where the building block consists of convolutional operations only, we added in Batch Normalization layers (*52*) and residual connections (*53*) for faster convergence and better performance. We also reduced the number of filters in each layer of the network by a factor of 8. The numbers of convolutional filters in the 4 stages of the encoding path are set to 8, 16, 32, and 64 respectively. The convolutional filter sizes were set to 3 × 3 across the network except for the kernels used for residual connection, which were set to 1 × 1. Our model ended up having 0.8 million trainable parameters. Given an input digital-mixing image, our model will output a pixel-wise classification (Fig. 5).

The model was trained with batches of images of size 160 × 160 pixels, randomly cropped from 240 × 240 images from our training set. The batch size was set to 20. The weights were optimized with the Adam optimizer against the categorical cross-entropy loss function:

$$E = -\frac{1}{r \cdot c} \sum_{r=1}^{h} \sum_{c=1}^{w} \sum_{k=1}^{5} \delta(y[r][c] = k) \cdot \log(y[r][c][k]), \quad [7]$$

where $h$ and $w$ represent the number of rows and columns in the image. $\delta$ is the indicator function, which evaluates to 1 if $y[r][c]$, true label of the pixel $(r, c)$, is $k$. This loss function takes the average of the negative log-likelihood for the target class across every pixel in one image. It penalizes the model when it infers a small probability for the target class.

The model was implemented using Python and TensorFlow (*54*). The training was performed on an NVIDIA GTX 1070 GPU with 8 GB memory. For the first dataset, approximately 1200 virus particles were cropped out manually, using FIJI (ImageJ). We used over half of the selected virus particles to synthesize 32, 240 × 240 test images as the test dataset. Each of these images contained 20 virus particles equally distributed among SARS-CoV-2, H1N1, HAdV and ZIKV. For the training dataset, each virus particle were either rotated or flipped before placed onto the background image. We generated (with repetition of particles) 500 images for training and 50 for validation. The neural network we used was similar to the one explained previously (main text Fig. 5), except that the number of trainable parameters is larger (Fig. S7A) to account for the addition of background signal in the dataset.

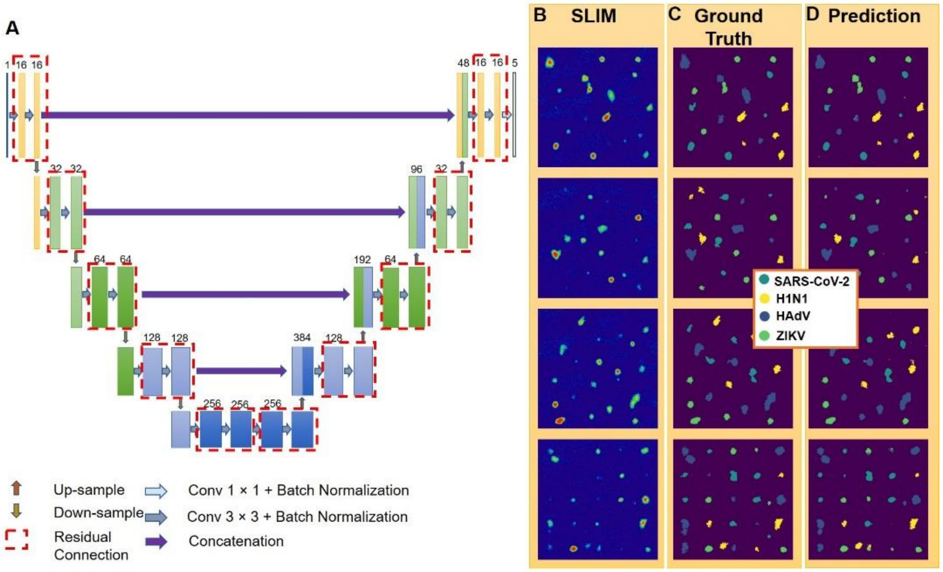

**Figure S7. Training a deep neural network to perform classification on the first digital-mixing dataset. A.** We used a modified version of U-Net for this semantic segmentation task. Besides reducing the number of parameters in the network to around 3 million, we also added in residual connection and batch normalization for faster convergence. **Model inference on images from the validation set and the test set. B.** Synthesized images of mixed virus particles. **C.** ground truth label. **D.** model inference.

Since pixel-wise segmentation accuracy did not reflect clearly the number of virus particles correctly and wrongly classified, we also introduced a particle-wise post-processing step (*55-57*). During the post-processing, we computed the average 5-class probability distribution across each virus particle:

$$p(R) = \frac{1}{|R|}\sum_{r \in R} p(r) \qquad [8]$$

$p(r) \in \mathbb{R}^{5\times1}$ is a vector that denotes the model's raw prediction for pixel $r$. Each entry in this vector gives the probability of pixel $r$ belonging to one of the 5 classes. Since our model used a softmax activation function as the output layer, the entries in $p(r)$ sums up to 1. $R$ represents a set of pixels that belong to the same virus particle. We enforced, via this post-processing strategy, the instance-level information onto the model's prediction such that the model will provide a unified label for one virus particle (Fig. S8).

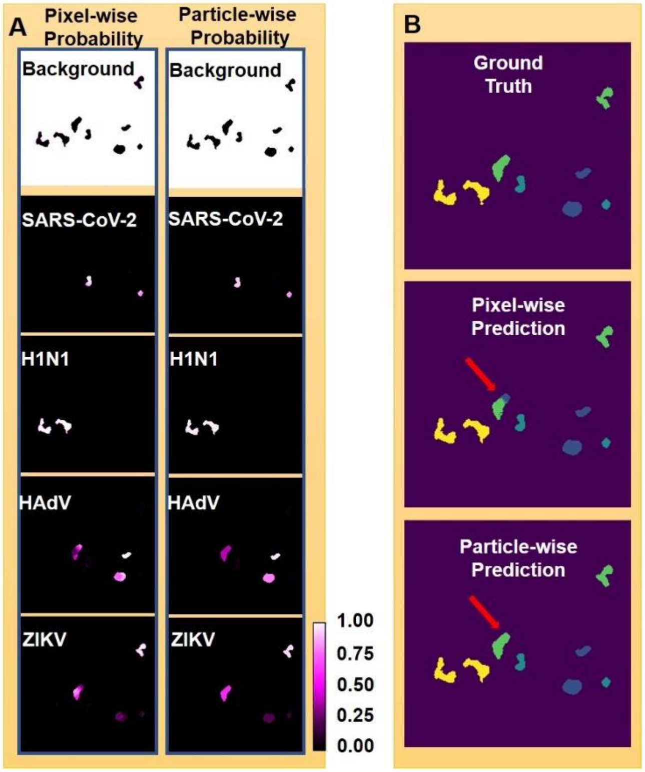

**Figure S8. Post-processing to enforce particle-level consistency. A.** To ensure all pixels in one virus particle has the same predicted label, we performed connected component analysis and averaged the probability distribution within each connected component. Left column: raw probability prediction; right column: probability distribution after post-processing. **B.** After post-processing, the predicted segmentation map no longer had different labels within one particle-region. This enabled us to compute, on an instance-level, the performance of our model.

### Network Performance

**First dataset:** SLIM images, ground truth and predictions are shown in Fig. S7 (B-D) respectively. After the application of post-processing (Fig. S8), we plotted the one-versus-all ROC curve and confusion matrix(Fig. S9). AUC values for four viruses are: 98% for SARS-

CoV-2, 98% for H1N1, 96% for HAdV and 97% for ZIKV (Fig. S9A). This model attained 0.80 precision and 0.88 recall for SARS-CoV-2, 0.82 precision and 0.73 recall for H1N1, 0.88 precision and 0.78 recall for HAdV, and 0.82 precision and 0.84 recall for ZIKV (Fig. S9B).

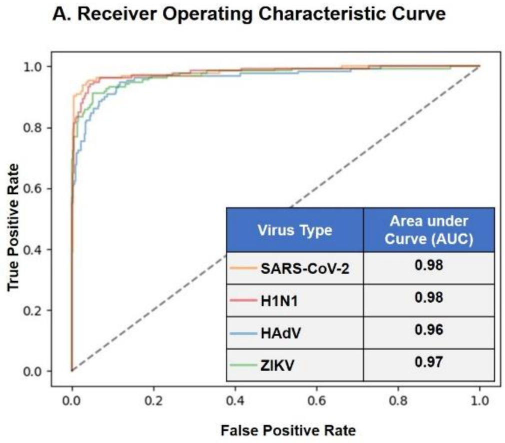

**B. Confusion Matrix**

| True Label \ Prediction | SARS-CoV-2 | Influenza-A (H1N1) | HAdV | ZIKV |
|---|---|---|---|---|
| SARS-CoV-2 (n = 133) | 87.97% | 11.28% | 0.75% | 0.00% |
| Influenza-A (H1N1) (n = 136) | 3.68% | 72.79% | 7.35% | 16.18% |
| HAdV (n = 130) | 12.31% | 3.08% | 78.46% | 6.15% |
| ZIKV (n = 134) | 9.70% | 3.73% | 2.99% | 83.58% |
| Average Precision | 0.80 | 0.82 | 0.88 | 0.82 |
| Average Recall | 0.88 | 0.73 | 0.78 | 0.84 |

**Figure S9. Model performance on the first test dataset (consisting of 32 240 × 240 images). A.** The receiver operating characteristic (ROC) curve of the model on the test dataset. The model achieved over 0.96 area-under-curve (AUC) for all 4 virus types on the test dataset. The area-under-curve (AUC) for each class is computed by setting that class as label 1 and all other class labels (background and the 3 remaining virus types) as label 0. **B.** The confusion matrix of the model inference on the test dataset. Each row represents the ground truth label while each column represents the prediction. For visualization purposes, each entry in the confusion matrix was normalized with respect to the number of true labels (sum of each row). The precision, recall are averaged across all images in the test dataset. Both the ROC curve and the confusion matrix are evaluated on a per-particle level.

**Second dataset**: The model was trained for 300 epochs. We set the initial learning rate to be 8e-5 and gave the model a warm up period (*58*) of 5 epochs. During the first 5 epochs, the learning rate increased linearly from 0 to 8e-5. After the warm up period, we implemented a simplified version of the cosine annealing strategy (*59, 60*) and gradually turned down the learning rate to 0 for better convergence. This model had a categorical cross-entropy loss of 0.03 on the validation dataset after 300 epochs of training. The model weights that gave the smallest loss value on the validation dataset were selected as our end model and used for evaluation.

Our model was evaluated on a test dataset consisting of 564 unseen images. Each of these images contained 8 - 32 virus particles randomly placed on a 240 × 240 blank image.

The same post-processing step as used for the first dataset was used for this dataset too and was implemented using the connected component analysis tool from scikit-image (*61*). The model inference achieved on average 0.8 precision and 0.85 recall for SARS-CoV-2 particles in the test dataset after post-processing (Fig. S10).

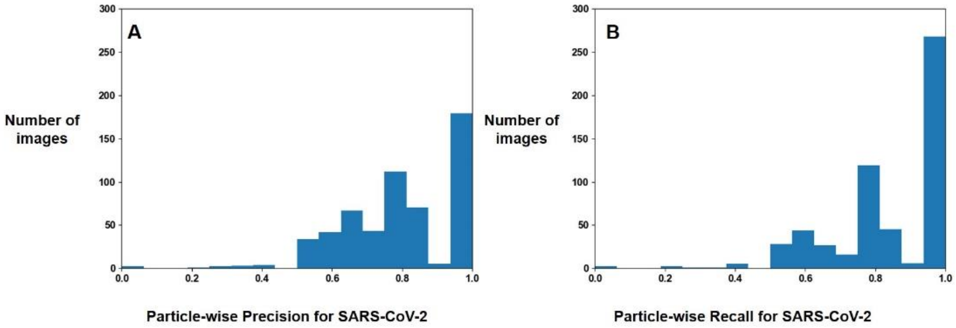

**Figure S10. Model Performance for SARS-CoV-2 with post-processing on the second dataset. A**. Histogram of particle-wise precision for SARS-CoV-2 evaluated on all 564 images in the test dataset. The average precision is 0.80. **B.** Histogram of particle-wise recall for SARS-CoV-2 evaluated on all 564 images in the test dataset. The average recall is 0.85.

All calculations are performed in Python using the scikit-learn library (*62*). Figure S11 shows the loss convergence for both our first (manual, Fig. S11A) and second (automatic, Fig.S11B) datasets.

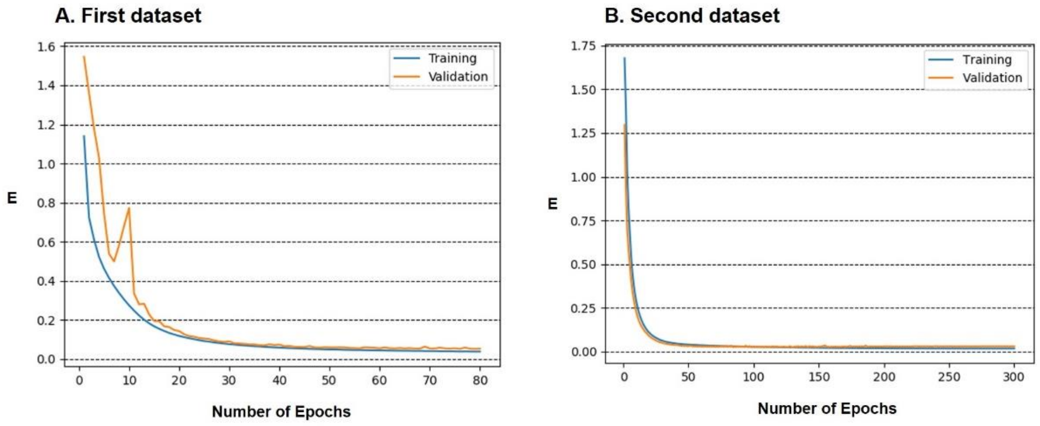

**Figure S11. Learning Curve Plot. A.** The learning curve plot of our model developed for the first dataset. **B.** The learning curve plot of our model developed for the second dataset. Both plots showed a good convergence between the validation loss and training loss of our models, indicating that our models did not underfit or overfit. E represents categorical cross-entropy Loss.
